# Morphogen gradients can convey position and time in growing tissues

**DOI:** 10.1101/2023.08.28.555133

**Authors:** Roman Vetter, Dagmar Iber

**Affiliations:** Department of Biosystems Science and Engineering, ETH Zürich, Schanzenstrasse 44, 4056 Basel, Switzerland; Swiss Institute of Bioinformatics, Schanzenstrasse 44, 4056 Basel, Switzerland

**Keywords:** tissue patterning, morphogen gradient, time, opposing gradients, neural tube, Sonic hedgehog

## Abstract

During embryonic development, cells coordinate fate decisions based on both position and time. While morphogen gradients have long been recognized as a source of positional information, how timing is synchronized across developing tissues remains unclear. We propose a simple mechanism by which morphogen dynamics can also encode time. If the morphogen source expands with a uniformly growing tissue and the gradient maintains a constant decay length, cells experience transient, hump-shaped signals that convey timing cues. Moreover, when two opposing exponential gradients with equal lengths—such as those in the vertebrate neural tube—interact, their product forms a uniform signal that can synchronize fate decisions across the entire tissue. With increasing gradient amplitudes, cells encounter a transient signal; with constant amplitudes, the signal decays—a feature of a depletion timer. This mechanism provides a generalizable principle by which morphogen gradients may coordinate both spatial and temporal aspects of patterning during development.

## 1 Introduction

During embryonic tissue development, graded profiles of signaling molecules, called morphogen gradients, guide spatial patterning and cell differentiation. The most basic model for this process is the French flag model, developed by Wolpert in the 1960s [1], according to which concentration thresholds define the locations of patterning domain boundaries in the tissue. In addition to their relative position, cells need to track the developmental stage to make the correct fate decisions in a timely fashion [2], such that the developmental program proceeds to build functional organs. A wide range of timer mechanisms have been identified [3, 4], including organism-wide timers based on temperature [5], nutrition [6], or hormonal changes [7], and local timers based on intracellular oscillators [8–10], accumulation or depletion processes [11–17]. However, how the various developmental processes and events are coordinated and synchronized across morphological fields remains largely unknown. We now propose a second developmental function of morphogen gradients: In addition to their well-established role as carriers of positional information, we show that morphogen gradients can convey also time during development under the right conditions.

The development of the central nervous system (CNS) has been established as a paradigm for gradient-controlled cell differentiation [18], and we will draw on the quantitative data that has been gathered for this model system. The CNS develops from the neural tube (NT) that runs along the rostral-caudal axis of the embryo. Neural development is controlled by opposing Sonic Hedgehog (SHH) and Bone morphogenetic protein (BMP) gradients that define different progenitor domains along the dorsal-ventral (DV) axis (Fig. 1). Quantitative analysis of the mouse NT at the forelimb level revealed that cell differentiation commences about two days after the start of NT development along the entire DV axis [19]. How this timepoint is set is unknown, but it is preceded by a transient response of the SHH and BMP pathways [20–24].

**Figure 1:**
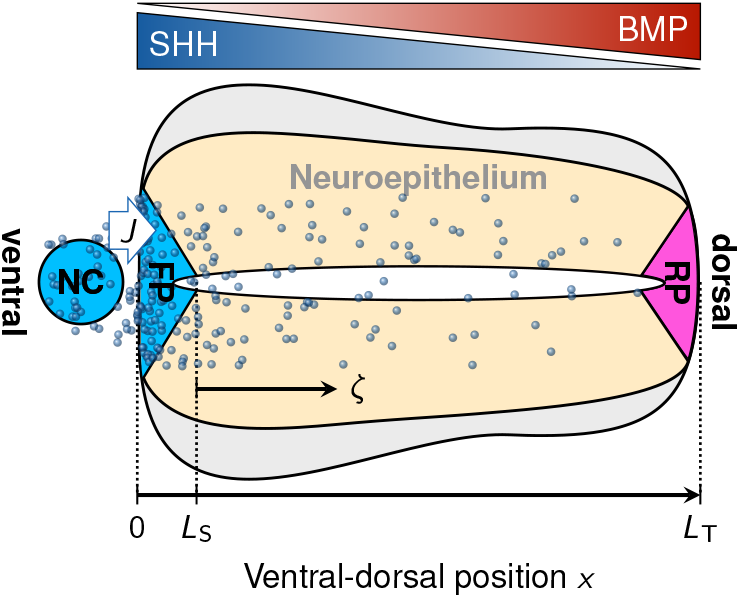
Patterning of the developing neural tube. Schematic cross section of the neural tube with floor plate (FP), roof plate (RP), and notochord (NC). SHH (blue molecules) is secreted from the NC and FP, and diffuses dorsally along the neuroepithelium, forming an exponential gradient. BMP is secreted from the RP, forming an anti-parallel gradient. The (positive or negative) net flux of SHH from the NC to the FP is represented by *J*. *ζ* = (*x* − *L*_S_)*/L*_T_ measures the relative position in the tissue, where *L*_S_ is the length of the morphogen source domain (the FP for SHH in the NT) and *L*_T_ the total tissue length (the ventral-dorsal size of the NT).

SHH and BMP are secreted from the two opposite ends of the NT. The SHH gradient and the SHH reporter GBS-GFP as well as the BMP readout pSMAD have been quantified and follow an exponential function in the patterning domain between floor plate and roof plate [22, 24]. While the gradient decay lengths are constant, the SHH gradient amplitude rises continuously over time [22]. Despite this gradual increase, the response to SHH, as monitored by GBS-GFP, is a transient hump, sharply rising at first and slowly receding later on [20–22]. The mechanisms that may produce such a response are elusive [22].

By solving a reaction-diffusion model, we demonstrate that the widening of the morphogen source in a growing tissue, as has been reported for SHH in the developing NT [19], results in a gradual increase in the morphogen amplitude, resembling that reported for SHH. In case of a constant gradient decay length, as is also the case for SHH in the NT, this rising amplitude results in hump-shaped temporal concentration profiles at fixed relative positions in the tissue. While the local concentration can convey positional information, the duration of the hump in the morphogen response can convey time. We further show that antiparallel exponential steady-state morphogen gradients can give rise to two types of timers that may synchronize developmental decisions along the expanding patterning axis.

## 2 Results

### 2.1 Morphogen concentration gradient

In systems with primarily diffusive morphogen transport, the morphogen gradient shape typically emerges as the solution of a reaction-diffusion equation. Diffusion at rate *D* and linear degradation or turnover at rate *k* give rise to exponential gradients of the form *c*(*x*) ∝ exp[−*x/λ*] with characteristic length 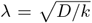, as derived further below. In the NT, the measured constant value of *λ* suggests that the spatial SHH gradient is in quasi-steady state on the patterning time scale [25], despite the time-dependent amplitude. This is consistent with measurements of the Hh dynamics in the *Drosophila* wing disc, where the Hh-GFP diffusion coefficient was determined as *D*_Hh_ = 0.033 ± 0.006 µm^2^ s^−1^ and the Hh-GFP turnover rate as *k*_Hh_ = 6.7×10^−4^ s^−1^ [26]. The characteristic time to steady state [27], *τ* = (1 + *x/λ*)*/*2*k*_Hh_ *≈* 25 minutes at *x* = *λ*, is thus short relative to the duration of NT patterning and expansion. The corresponding gradient decay length 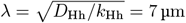 of Hh-GFP in the *Drosophila* wing disc is remarkably close to that measured for SHH-GFP, *λ ≈* 13 µm, in the mouse NT [24, 28], and somewhat shorter than *λ ≈* 20 µm for untagged SHH [22]. The difference between Hh-GFP in the *Drosophila* wing disc and SHH-GFP in the mouse neural tube could be due to differences in the regulatory interactions [28, 29]. In conclusion, we will now assume that the gradient is in quasi-steady state on the growing domain due to the separation of timescales.

We approximate the tissue by a 1D domain of total length *L*_T_, with the coordinate *x* running from the side of the source, *x* = 0, to the other end, *x* = *L*_T_. In case of SHH in the NT, the domain would start at the ventral limit of the floor plate (FP), and extend to the dorsal end of the roof plate (RP) (Fig. 1). The morphogen distribution along the uniformly expanding domain can then be described with a 1D steady-state reaction-diffusion equation [30]:

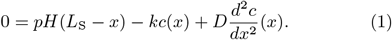

Here, *c* denotes the morphogen concentration, *p* the morphogen production rate, *k* the turnover rate, *D* the diffusion coefficient, and *H* the Heaviside step function that restricts the morphogen production to the source of length *L*_S_:

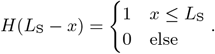

We assume that the boundary far from the source is impermeable, and use a zero-flux boundary condition at *x* = *L*_T_:

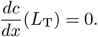

We will show later that this assumption has little impact on the predicted gradient shape.

In the NT, there is no obvious diffusion barrier at the opposite boundary, as SHH diffuses between the adjacent SHH-secreting notochord (NC) and the FP. Accordingly, we explore the impact of SHH diffusion across this boundary with a finite morphogen flux *J* in or out of the NT (Fig. 1):

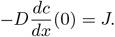

Eq. 1 can be solved with basic calculus by splitting the unknown concentration profile into two domains, one containing the morphogen source, and another one in which the patterning takes place:

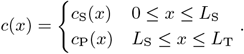

At the interface between source and patterning domain (*x* = *L*_S_), we require continuity in the concentration and flux:

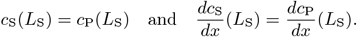

With these boundary conditions, the solution for the morphogen concentration reads

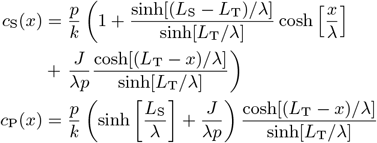

with gradient decay length 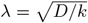. Note that the boundary flux appears only in the dimensionless ratio *J*/*λp*.

The position where the zero-flux boundary condition is imposed opposite of the morphogen source has a negligible quantitative impact on the concentration gradient, as long as it is far away from the source (*L*_T_ ≫ *L*_S_), which is always the case in the NT [19]. One can thus make a simplification without altering the solution in any practically measurable way. Instead of imposing *dc/dx* = 0 at *x* = *L*_T_, we can impose it infinitely far away from the source at *x* = *∞*. With this modification, the concentration simplifies to a pure exponential in the patterning domain:

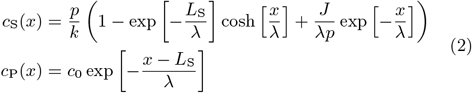

where *c*_0_ = *c*(*L*_S_) is the the amplitude to the exponential gradient in the patterned tissue, given by

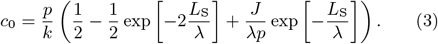

Importantly, assuming constant kinetic parameters *p/k, J/λp* and *λ*, the signaling amplitude increases with an expanding source size (Fig. 2A).

**Figure 2:**
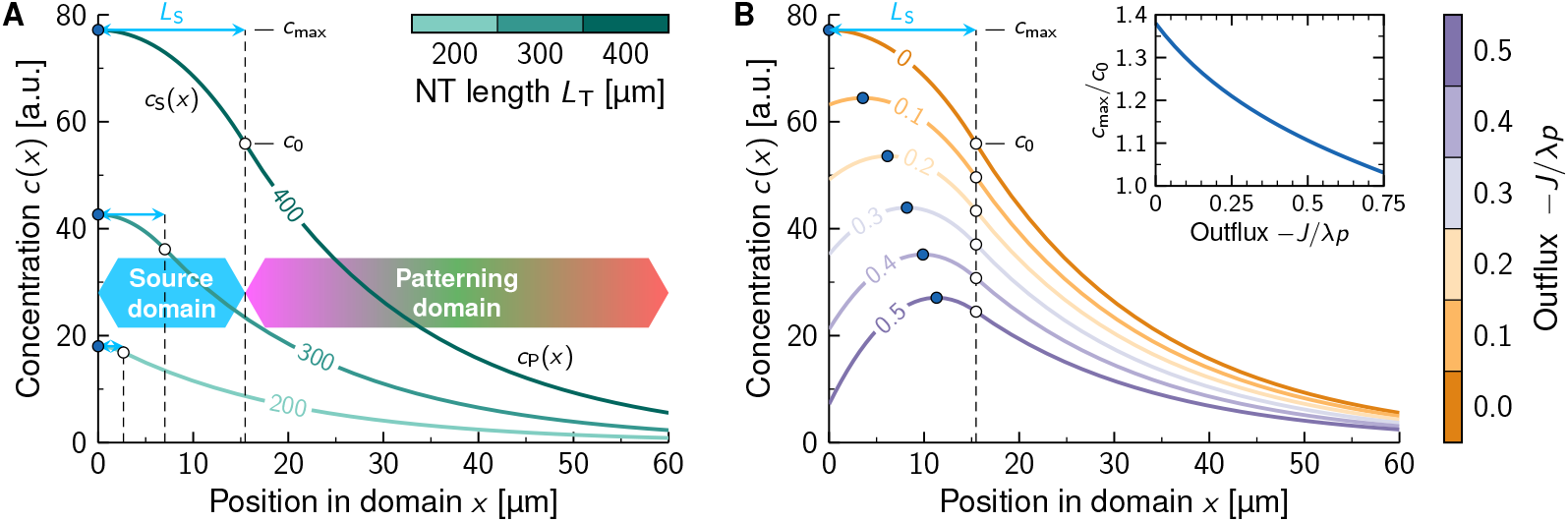
SHH gradient in the growing neural tube. **A**, Eq. 2 is plotted for different NT lengths using the quadratic fit *c*_max_(*L*_T_) from Fig. 3A, for the case without flux (*J* = 0). **B**, Eq. 2 at fixed total NT length *L*_T_ = 400 µm for different morphogen outflux values from the FP to the NC, as labeled. Maximum concentrations *c*_max_ (blue dots) and boundary concentrations *c*_0_ (white dots) are indicated. With increasing outflux, the maximum concentration approaches the source boundary amplitude: *c*_max_ *→ c*_0_ (B, inset).

For a positive influx *J* > 0, the maximum concentration occurs at *x* = 0. A negative flux *J* < 0 (corresponding to an outflux of SHH from the FP to the NC), on the other hand, shifts the peak concentration to the interior of the source (Fig. 2B). The concentration profile obtained with a moderate outflux resembles the reported profiles of SHH in the NT [22] and Bicoid (Bcd) in the early *Drosophila* embryo [31]. The maximum can be found by requiring that *dc/dx* = 0. It is

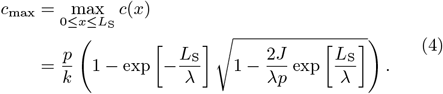

This maximum is attained at position

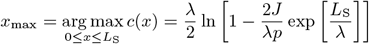

which progressively approaches the source-pattern boundary at *x* = *L*_S_ (the dorsal FP boundary in the NT) with increasing morphogen outflux. Accordingly, the maximum concentration *c*_max_ approaches the exponential amplitude *c*_0_ with increasing outflux (Fig. 2B, inset) and would reach it at *J*/*λp* = − sinh[*L*_S_*/λ*] if the concentration did not hit zero already at *J*/*λp* = exp[−*L*_S_*/λ*]−1. Inside the patterning domain, the exponential shape of the gradient is unaffected by *J*; only the amplitude is slightly shifted according to Eq. 3. To reduce the mathematical complexity in our further analysis below, we will therefore express the morphogen gradient in the patterning domain by

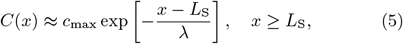

which is valid in good approximation, considering the reported shape of the Bcd and SHH profiles [22, 31]. The subsequent results are qualitatively unaffected by this simplification.

Note that a morphogen outflux is not the only possible explanation for the observed concentration peak inside the source domain. Other potential origins include non-uniform morphogen kinetics (production and degradation), non-uniform transport properties, or geometric effects of the wedge-shaped FP. The precise nature of the cause for the observed shape of *c*_S_ is unimportant for the following analysis.

### 2.2 Gradient amplitude dynamics

According to Eqs. 3 and 4, the gradient amplitude increases with the size of the source. The maximal SHH amplitude and the FP length have been reported for different NT lengths (Fig. 3A) [19, 21, 22]. We can now use these measurements to parameterize our model.

**Figure 3:**
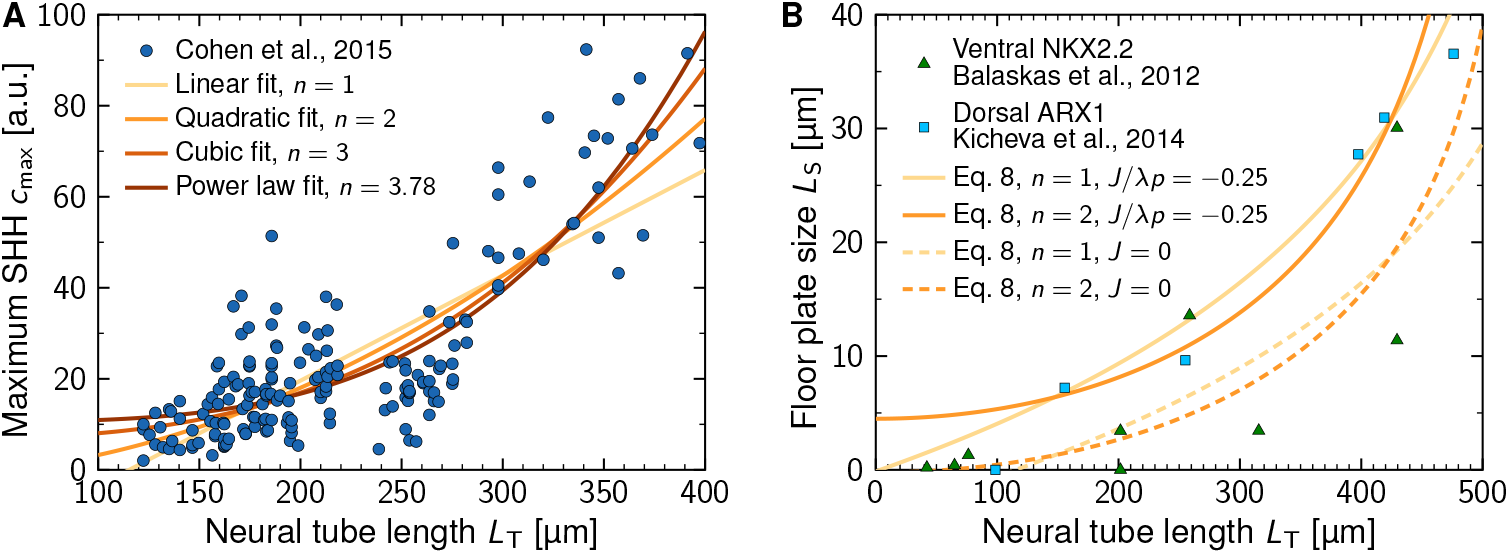
SHH gradient amplitude in the growing neural tube. **A**, Polynomial models (colored lines) fitted to the increasing SHH gradient amplitude (blue dots, extracted from Ref. [22]) in the growing NT. **B**, The floor plate size inferred from the fitted maximum SHH concentration using Eq. 7 matches the reported data from Refs. [19, 21] within a moderate range of outflux values (solid and dashed lines). The inferred flux values (Table 1) lie in the shown range, close to zero. Developmental time was converted to NT length using the linear expansion law by Cohen et al. [22].

In previous work [22], a linear model (*n* = 1) was fitted to the maximum SHH concentrations. While this fits the data reasonably well, the fit improves for higher-order polynomial functions,

**Table 1:** SHH gradient amplitude dynamics. Parameters were obtained by least-squares fitting Eqs. 6 and 7 to the data in Fig. 3. RMSE and adjusted *R*^2^ values refer to the concentration fit in Fig. 3A.

| Model | Parameter | Unit | Value | SE | RMSE [a.u.] | Adjusted $R^2$ |
| --- | --- | --- | --- | --- | --- | --- |
| linear ($n = 1$) | $p/k$ | a.u. | 115 | 29 | 11.8 | 0.616 |
| | $J/\lambda p$ | — | -0.02 | 0.15 | | |
| | $\alpha$ | — | -0.232 | 0.027 | | |
| | $\beta$ | $\mu\text{m}$ | 497 | 29 | | |
| quadratic ($n = 2$) | $p/k$ | a.u. | 140 | 30 | 10.9 | 0.674 |
| | $J/\lambda p$ | — | -0.01 | 0.14 | | |
| | $\alpha$ | — | -0.012 | 0.011 | | |
| | $\beta$ | $\mu\text{m}$ | 533 | 14 | | |
| cubic ($n = 3$) | $p/k$ | a.u. | 183 | 33 | 10.3 | 0.706 |
| | $J/\lambda p$ | — | -0.04 | 0.13 | | |
| | $\alpha$ | — | 0.037 | 0.006 | | |
| | $\beta$ | $\mu\text{m}$ | 524 | 8 | | |
| power law ($n$ free) | $p/k$ | a.u. | 228 | 38 | 10.3 | 0.711 |
| | $J/\lambda p$ | — | -0.07 | 0.12 | | |
| | $\alpha$ | — | 0.046 | 0.008 | | |
| | $\beta$ | $\mu\text{m}$ | 518 | 19 | | |
| | $n$ | — | 3.78 | 0.40 | | |

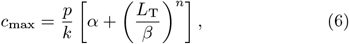

and is optimal for *n* = 3.78 ± 0.40 (Fig. 3A, Table 1). However, since the variability in the data is too large to permit a conclusive identification of the rate at which the maximum SHH concentration increases, we maintain *n* as a free parameter and discuss particular values in the following. Balancing Eq. 6 with Eq. 4, we can infer how the FP must expand with the NT to yield the measured increase in the SHH amplitude. The resulting relationship

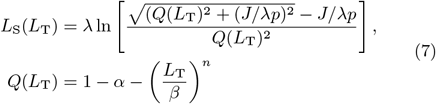

matches the reported FP length for a range of net SHH outflux values of about −*J*/*λp* ∈ [0, 0.25] (Fig. 3B), although a statistical zero is the optimal flux value we inferred with the sparse available data (Table 1). Even though the FP expands passively with the uniformly growing domain [19], the functional relationship is non-linear because the FP is not present from the start, but is only induced by SHH from the notochord. This is in agreement with a recent model of FP induction that suggested that *J ≈* 0 after early establishment, followed by tissue growth and SHH production in the FP [32].

Note that the prefactor *p/k* merely sets the arbitrary concentration scale. By fitting Eq. 6 alone to measured amplitudes, the value of *p/k* is not uniquely defined and this prefactor can be accommodated in the values of *α* and *β* with appropriate unit adjustment. With the simultaneous fit of Eq. 7 to measured source lengths, however, one can determine optimal values for *p/k, J/λp, α* and *β* uniquely. Nevertheless, *p/k* still only represents the arbitrary concentration scale used in the SHH amplitude data [22].

### 2.3 Hump-shaped gradient dynamics on growing domains

In a uniformly growing tissue, stationary cells remain at a fixed relative position *ζ* ∈ [0, 1]. What is the signal they are then exposed to over time? Combining Eqs. 5 and 6, the morphogen concentration can be evaluated at relative positions *ζ* = (*x* − *L*_S_)*/L*_T_ along the patterning domain (Fig. 1), and at developmental stages of tissue growth as quantified by the total domain length *L*_T_:

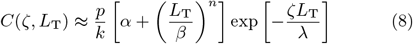

At fixed *ζ*, the concentration first increases as the tissue expands, before it starts declining again—a biphasic (hump-shaped) transient signaling profile (Fig. 4A). For the quadratic relationship (*n* = 2), the reversal point occurs earlier than for the linear model (*n* = 1) (Fig. 4B). As the NT grows linearly in time [19], the NT length where the peak morphogen concentration is attained, *L*_peak_, directly relates to developmental time, allowing the SHH dynamics to be compared to the reported signaling dynamics [21, 24]. The measured SHH gradient decay length *λ* and amplitude (*α, β, n*) directly determine the NT length at which the peak *C*_peak_(*ζ*) = *C*(*ζ, L*_peak_) is reached at a given relative position *ζ*. This relationship can be found from *∂C/∂L*_T_ = 0 and it reads

**Figure 4:**
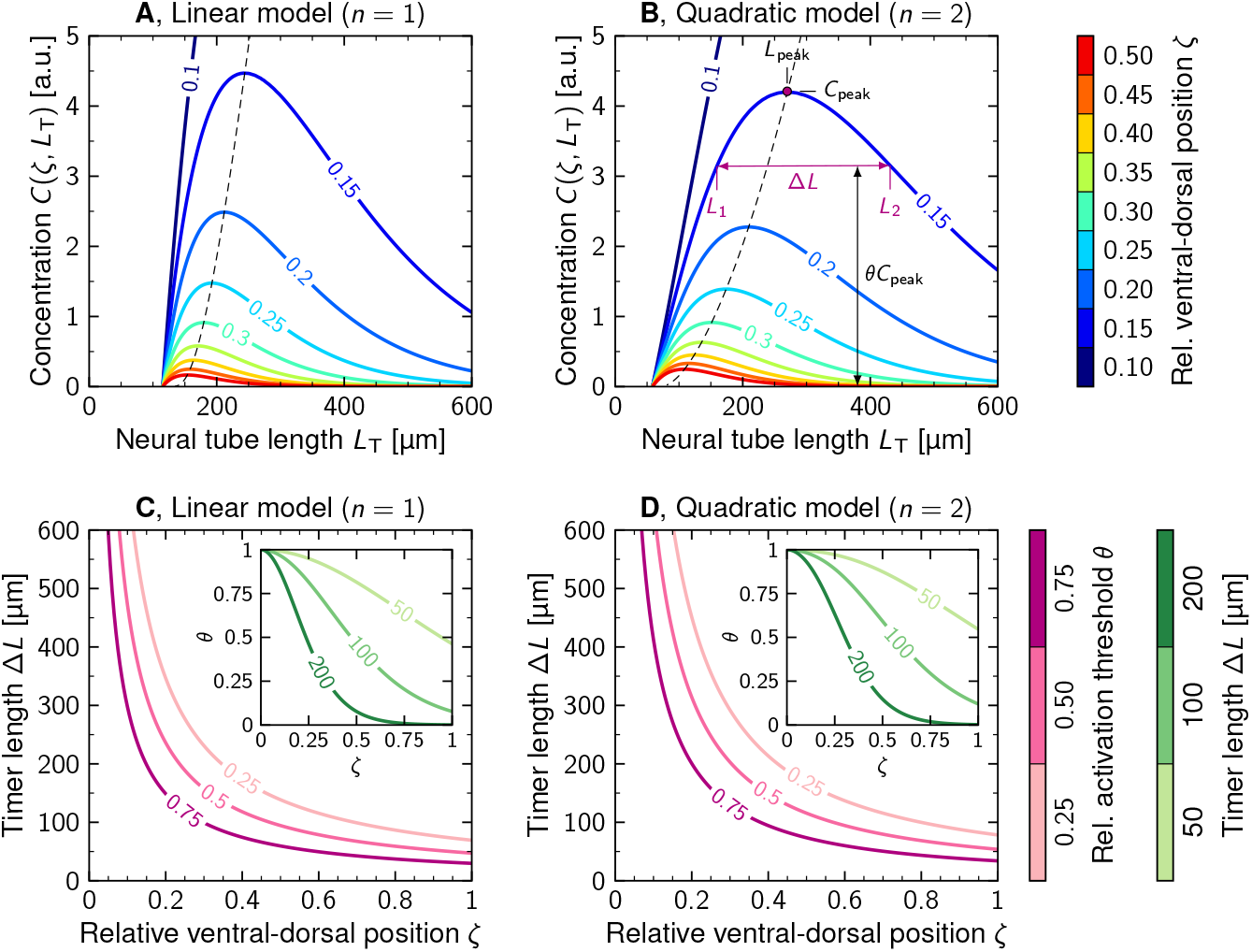
Transient gradient dynamics on growing domains. Time-dependent concentration at different ventral-dorsal positions *ζ* (colors) for a linearly increasing gradient amplitude (**A**) and a quadratically increasing gradient amplitude (**B**). Eq. 8 is plotted using the parameters from the measured SHH gradient (Table 1). The dashed lines show the evolution of the peak concentration *C*_peak_, given by Eqs. 10 and 11. Duration (in terms of domain expansion Δ*L* over time) of the transient activation of the SHH pathway through exceedance of a fraction *θ* of the peak concentration, as a function of the relative position in the pattern, *ζ*, for a linearly increasing gradient amplitude (**C**, Eq. 12) and a quadratically increasing gradient amplitude (**D**). Insets show the inverse relationship, the relative activation threshold as a function of position for fixed timer lengths.

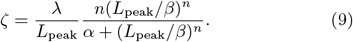

In case of the linear fit (*n* = 1), the NT length where the peak is attained follows as

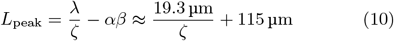

whereas it is

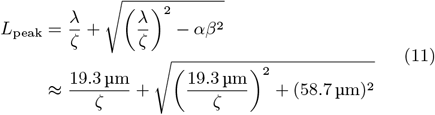

for the quadratic fit (*n* = 2). The peak is thus attained earlier further away from the source (Fig. 4A,B). Generally, the condition for a hump to appear is that

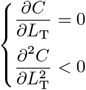

has a solution within the range of *L*_T_ as the tissue expands^1^. For a constant *λ* and *c*_0_(*L*_T_) *≈ c*_max_(*L*_T_) as in the mouse NT, this reduces to the condition that Eq. 9 has a solution for *L*_peak_ in the range of physiological tissue lengths. For general *n*, this cannot be reduced further analytically, but for *n* = 1, 2, such that the gradient amplitude rises enough at short domain lengths, a hump indeed forms.

The local hump corresponds to a transient activation of the SHH pathway. At fixed relative positions *ζ* in the patterning domain, the local morphogen concentration crosses any threshold concentration below the peak twice: Once to exceed it, and a second time at a later timepoint to drop below it again. For any threshold *θ* ∈ (0, 1), the duration of the transient activation can be determined numerically as the distance Δ*L* = *L*_2_ − *L*_1_ between the two crossing points where *C*(*ζ, L*_1_) = *C*(*ζ, L*_2_) = *θC*_peak_(*ζ*) (Fig. 4B). Δ*L* corresponds to the length of the growing domain gained over the duration of signaling, which we take as a proxy for the developmental stage here, to make our results independent of the tissue growth rate. For a linearly increasing morphogen gradient amplitude (*n* = 1), it can be found by inverting Eq. 8, and it reads

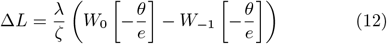

independent of *α* and *β*. Here, *W*_*k*_ denotes the *k*-th branch of Lambert’s *W* function (the product logarithm), and *e* is Euler’s constant. The timer duration is thus inversely proportional to the relative position in the patterning domain, Δ*L ~ ζ*^−1^ for *n* = 1 (Fig. 4C). For a quadratically increasing morphogen gradient amplitude (*n* = 2), it can be found numerically, and still approximately follows *~ λ/ζ* curves (Fig. 4D). Supposing a uniform threshold *θ* in the domain, the transient activation of the SHH signaling pathway is thus longer close to the source and shorter further away from it. Conversely, if instead of the activation threshold *θ*, the timer length is to be uniform across the domain, then the activation threshold needs to drop in a nonlinear fashion with increasing distance from the source (Fig. 4C,D, insets), similarly to the French flag model for patterning in space.

We conclude that a hump-shaped local morphogen profile can arise if the morphogen gradient amplitude can be related to the patterning domain length via a polynomial function of order two or smaller. For a cubic (*n* = 3) or higher-order relationship, transient responses occur only close to the source. Further out, the concentration declines continuously. The gradient parameters directly determine the timing of the hump; there are no free parameters.

### 2.4 Synchronized timing by opposing morphogen gradients on growing domains

During NT development, cell fate decisions are largely synchronized along the patterning axis [19, 34, 35]. How such a synchronization can be achieved across a large, growing morphogenetic field, and how timing is tied to developmental progress, is still largely elusive. While a single gradient can coordinate timing along the patterning axis, synchronization would require a finely tuned distance-dependent readout threshold (Fig. 4C,D). We now show that opposing steady-state morphogen gradients with equal decay length, as found in the vertebrate neural tube [24], can serve as developmental timers that synchronously trigger a fate switch once the tissue reaches a critical size.

Opposing exponential gradients can be written as

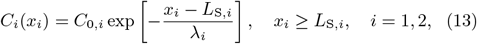

where *x*_*i*_ denotes the distance from the tissue boundary with the respective morphogen source. On a domain of total length *L*_T_, *x*_1_ = *x* for one gradient and *x*_2_ = *L*_T_ − *x* for the second, anti-parallel gradient. If both gradients have an equal decay lengths (*λ*_1_ = *λ*_2_), but possibly different amplitudes *C*_0,1_ and *C*_0,2_, the product of their concentrations,

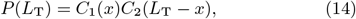

is constant along the entire patterning axis, and declines uniformly in the growing tissue, as required for a depletion timer (Fig. 5A,B). The combined concentration of the opposing gradients can time development if the domain length *L*_T_ changes over time. The timer length is then given by the amount of tissue expansion required to let the product concentration drop below a threshold, *P* = *θC*_0,1_*C*_0,2_ (Fig. 5C):

**Figure 5:**
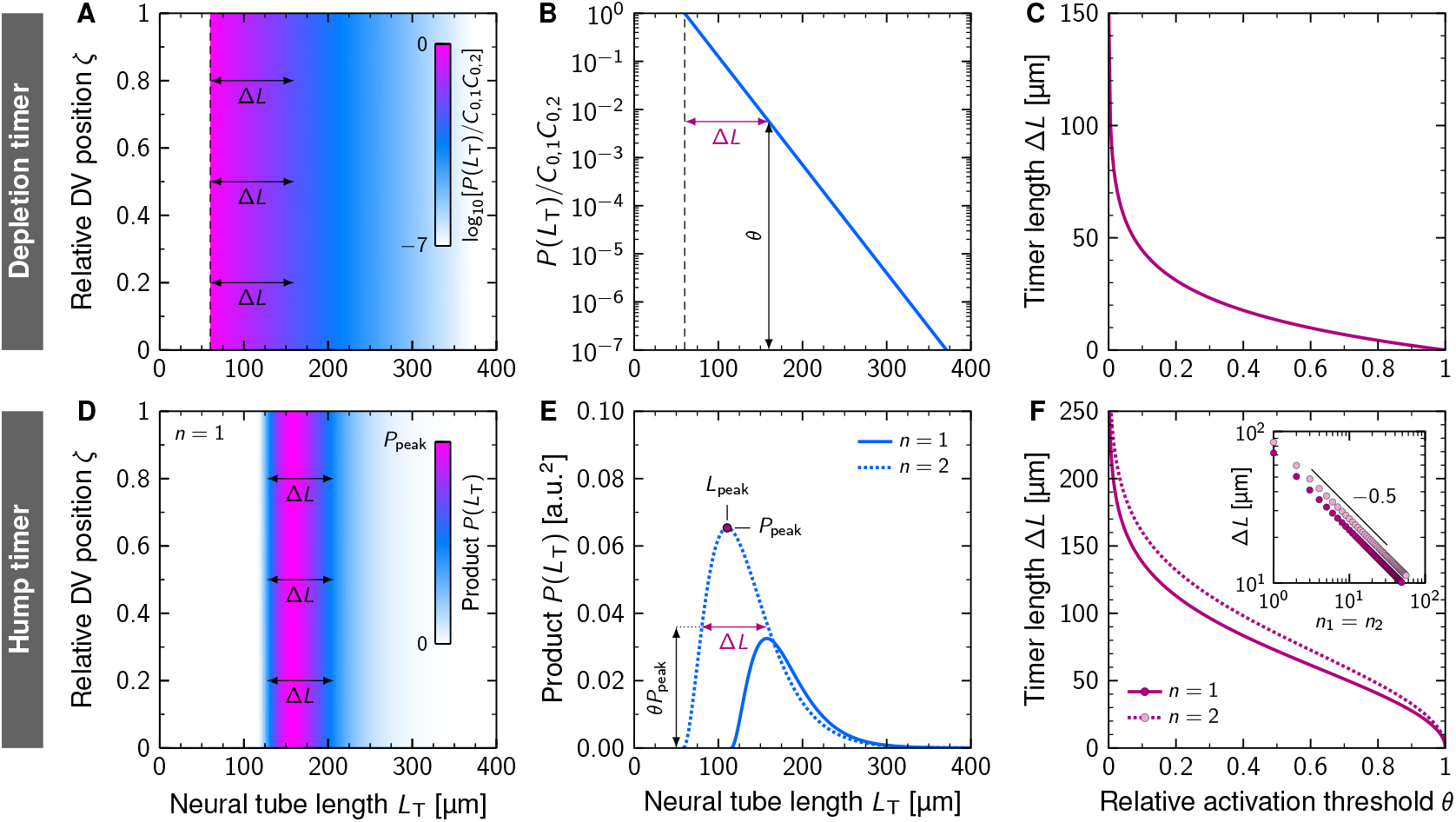
synchronized timing by opposing morphogen gradients on growing domains. **A**, Time-dependent product of concentrations of anti-parallel morphogen gradients. Eq. 14 is plotted for fixed gradient amplitudes and source sizes, resulting in a depletion timer that is synchronous everywhere in the pattern. Note the logarithmic color scale. **B**, The product concentration drops below a threshold (i.e., depletes) after the tissue has grown by Δ*L*. **C**, Length of the depletion timer as a function of the sensing threshold. **D**, Time-dependent product of concentrations of anti-parallel morphogen gradients for increasing amplitudes as measured for the SHH gradient (*n* = 1, Table 1), assuming that the opposing BMP gradient is mirror-symmetric to it. Note the linear color scale. **E**, Time evolution of the product concentration for *n* = 1, 2, showing a transient hump as the NT lengthens over time. **F**, Length of the timer as a function of the activation threshold (relative to the peak signal for non-dimensionalization), as indicated in **E**. Inset: Scaling of the timer length with the Hill coefficients *n*_1_ = *n*_2_ for a transcriptional implementation of the hump timer, at fixed *θ* = 0.5.

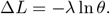

If, on the other hand, the amplitudes rise over time, as in the vertebrate NT, then the transient product concentration exhibits a synchronous hump. Using the gradient amplitude fit for SHH in the mouse NT for both the SHH and the BMP gradients (*C*_0,1_ = *C*_0,2_ = *c*_max_, Eq. 6), the peak *P*_peak_ = *P* (*L*_peak_) occurs at a critical tissue size of *L*_peak_ *≈* 158 µm for *n* = 1 and at *L*_peak_ *≈* 110 µm for *n* = 2, everywhere in the patterning domain. Note that *L*_peak_ would be different if the amplitude of the opposing gradients differed. Before the critical size is reached, the morphogen product increases uniformly in the entire tissue, and depletes uniformly afterwards (Fig. 5D,E). As with just one morphogen gradient, the duration of the timer with opposing gradients, measured in terms of tissue expansion Δ*L*, depends on the activation threshold above which the timer is turned on. The lower the threshold, the longer the timer, with uniform timer length and activation threshold across the entire pattern (Fig. 5F). (Note that the same argument holds with an absolute threshold *P*_*θ*_ instead of a relative threshold *θ*. Cells do not need to know *P*_peak_.) Time-dependency in the amplitudes will thus affect the exact timing, but not the synchronization along the patterning axis. All that is required for synchronization is that both morphogen gradients have equal *λ*, i.e., equal *D/k*.

Such a product readout could be implemented biologically for example through a transcriptional gene regulatory network that links both ligands through a logical AND gate [36]. Assuming Hill–Langmuir kinetics, this would result in an activation function of the form

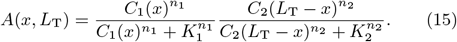

The sufficient conditions for the activation to be uniform across the entire patterning domain (*A*(*x, L*_T_) ≡ *A*(*L*_T_)), and thus for synchronous timing, are then that both Hill functions are in the linear regime (i.e., *K*_*i*_ ≫ *C*_*i*_) and

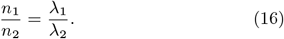

Notably, this transcriptional timer form allows to relax the constraint on equal gradient lengths through corresponding adjustment of the Hill coefficients *n*_1_, *n*_2_.

A depletion timer based on this readout that is triggered when the activation drops below the threshold *A*_*θ*_ would then have a timer duration (again measured by the domain elongation Δ*L*) of

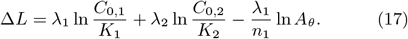

For this length to be positive, the activation threshold needs to be sufficiently small:

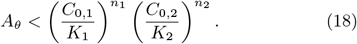

A hump timer based on this readout that is started when the activation first exceeds *A*_*θ*_ and stopped again when it drops back below *A*_*θ*_ has a duration Δ*L* = *L*_2_ − *L*_1_ that can be found numerically from the difference of the two solutions to *A*(*L*_1,2_) = *A*_*θ*_ for a sufficiently low threshold *A*_*θ*_. In relative terms with *A*_*θ*_ = *θA*(*L*_peak_), the duration Δ*L*(*θ*) behaves qualitatively as shown in Fig. 5F, but shortens with higher Hill coefficients: 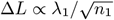 at fixed relative thresholds *θ* (Fig. 5F, inset).

## 3 Discussion

We have shown that the passive expansion of the morphogen source with a uniformly expanding developing tissue results in transient, hump-like morphogen kinetics along the patterning domain. Opposing gradients with equal decay length, as found in the neural tube [24], allow for the synchronization along the entire pattern. This new patterning paradigm offers a simple mechanism for the simultaneous control of position and timing by the morphogen gradient(s) alone. With such a mechanism, the position of progenitor domain boundaries can be controlled by the gradient amplitude and corresponding readout thresholds, while the timing of developmental processes can, in principle, be controlled with a readout of the transient nature of the concentration profiles. An expanding source and opposing gradients are found in many growing patterning systems, such that this could constitute a general timer paradigm.

Note that the hump-shaped local morphogen concentration inside the expanding patterning domain is not a dilution effect. Our analysis is based on a steady-state analysis of morphogen gradients, and requires only three main ingredients: 1) an increasing amplitude of the morphogen gradient, 2) a (local or global) expansion of the patterned tissue, and 3) a non-scaling (e.g., constant) gradient decay length. These conditions are met in the developing neural tube, for example, where the morphogen source domain co-expands with the uniformly growing tissue, while the gradients maintain their *λ*. In such tissues, the local hump that may allow cells to measure developmental time emerges naturally from uniform tissue expansion. A cell situated at a fixed relative position in the tissue effectively moves away from the morphogen source. There are thus two counter-acting effects at play: A rising signal from the growing gradient amplitude, and a signaling decline from the passive cell movement away from the source, down the gradient. Note that uniform growth is not absolutely required, but it is the simplest scenario that can give rise to a hump timer. The minimal requirement is that the portion of the tissue in which development is to be timed move away from the source, down the morphogen gradient. In the case of uniform growth, as observed in the neural tube, this will thus work for the entire domain. In case of non-uniform growth, the paradigm can apply only to those regions that experience a net expansion of the tissue between it and the morphogen source.

A key advantage of gradient-based timers is the spatial control and alignment of cell populations over large distances. For intracellular timers, entrainment mechanisms are required to align the response of neighboring cells [37]. In case of Notch signaling, these rely on local cell-to-cell interactions. Alternatively, the secretion of diffusible factors can coordinate local cell response, as proposed for TGF-*β* in hindbrain development [14]. For gradient-based timers, there is a firm bond between the rate of the gradient amplitude increase and the timing of developmental processes. As a result, spatial patterning and timing become intricately linked. The domain length at the morphogen peak is sensitive to changes in the gradient amplitude dynamics and the gradient decay length (Eqs. 10 and 11). This sensitivity offers a mechanism to adjust time and length scales during evolution. Human embryos are larger than mouse embryos and develop more slowly. If the longer half-life that has been reported for the human proteome [38] translates into a larger maximal gradient amplitude (Eq. 4) and gradient decay length, the gradient-based timer mechanism will not only delay the peak of morphogen signaling in human embryos, but will also shift it to larger domain lengths. Even if the production rate slows down in parallel to the decay rate due to a lower metabolic rate [39], a shift to longer domains would still be obtained. Gradient-based timers and those based on the local coupling of intracellular timers may thus have evolved in parallel to address different challenges in development. These are hypotheses that remain to be tested in the future.

Gradient-based timers can only yield precise patterning and timing if the positional error of their readout is sufficiently low across the entire patterning field. How far morphogen gradients are precise enough is a matter of scientific debate [24, 30, 40–42] and requires further clarification. In the mouse NT, the GBS-GFP and pSMAD profiles were imaged with 8-bit imaging depth [24], allowing only a 256-fold intensity range to be detected [30]. Beyond a theoretical maximum of 5.5*λ* = 110 µm from the morphogen source, these measurements therefore cannot contain any signal anymore, such that experimental data on the actual spatial range of gradients remains absent. Methodological advances will be required to shed more light from the experimental side onto this, considering also that absolute morphogen abundance in the NT has not been amenable to experimental quantification so far. Meanwhile, theoretical studies have found that for physiological noise levels, the positional error of morphogen gradients can be just a few cell diameters [33, 43] or even lower [44] across the entire NT, if copy numbers remain large enough also beyond the 5.5*λ* that have been experimentally covered. Experimental evidence in favor of the SHH gradient ranging far enough is that cells in the very dorsal part of the NT are sensitive to SHH from the FP [45]. Importantly, the SHH gradient amplitude is perfectly robust to alterations in gene dosage in that NT patterning is unchanged in *Shh* heterozygous mutants [46]. How this extraordinary robustness is obtained remains unknown, but it is also observed in other patterning systems such as the mouse limb [47]. Together, this suggests to us that the SHH gradient is likely sufficiently robust and precise to serve as developmental timer, but future research might help clarify this further.

In the mouse NT, transient SHH kinetics precede the onset of differentiation [19–22, 48, 49]. A transient response is obtained only with a linear or quadratic fit to the SHH amplitude data with a gradient-based timer, but not with the statistically favored higher order power-law fit (*n* = 3.78, Table 1). However, due to the spread in the available SHH amplitude and FP length data (Fig. 3), the optimality of *n* = 3.78 is likely not statistically robust, and growth laws other than the power law used here could be similarly adequate. Given the technical challenges in measuring SHH gradients and FP lengths in the NT, the measured SHH amplitude and/or FP length data could be inaccurate [30]. To clarify the exact functional form of the amplitude increase over developmental time, even denser measurements at the dorsal end of the floorplate might be useful. Measurements of the SHH flux *J* between the notochord and the FP could further help test the steady-state assumption in the NT. Our current fitting of the relatively sparse floorplate size data leaves room for a range of possible flux values.

We note that a timer based on the measured SHH gradient yields a transient response that is much slower and delayed compared to the measured kinetics of the SHH pathway [21, 22]. To achieve the same duration along the patterning axis, the readout threshold, *θ*, would need to decline with distance from the source (Fig. 4C,D). However, also in this case, the peak would still be reached too late close to the source (Fig. 4A,B). A timer based on the opposing SHH and BMP gradients would allow for the synchronization of cell differentiation along the entire patterning axis with a single readout threshold at a NT length consistent with the observed onset of differentiation.

A transient activation of the SHH pathway is observed also in cultured neural stem cells, and when the SHH pathway is stimulated permanently [14, 22, 38]. Models that include both diffusible morphogens and intracellular regulatory networks will be important to understand how tissues exploit their specific advantages to robustly coordinate cell fate decisions in space and time — and to what extent they are redundant. Important biological processes are often controlled by redundant mechanisms [50], a prominent example being hindbrain development. While TGF-*β* terminates the SHH response and thereby controls the concomitant switch in cell differentiation from motor neurons to serotonergic neurons (5HTN), the SHH response still terminates in the absence of TGF-*β* signaling, albeit somewhat delayed [14, 51].

We motivated our analysis with observations from the developing neural tube as it has the required properties, i.e, uniform tissue expansion, increasing amplitude, constant decay length, and two opposing gradients that can allow synchronization. Further experimental perturbations of the morphogen gradients will be necessary to test the proposed gradient timer mechanism in the NT, assess its role relative to intracellular timers, and its compatibility with regulatory interactions that are either downstream or affect the gradients themselves. Uniform domain expansion and the parallel expansion of the sources of opposing morphogen sources are observed also in many other developing tissues, but so far only few have been analyzed quantitatively [18]. In case of the Bicoid gradient in the *Drosophila* blastoderm, growth terminates before Bcd-dependent patterning [52] such that our novel timer paradigm cannot apply to the Bcd gradient. Temporal averaging [53], as explored in the syncytial early *Drosophila* embryo [54], and its potential effect on timing and synchronization in the gradient-based timer introduced here, could nevertheless be of interest in future work.

In case of Decapentaplegic (Dpp) in the *Drosophila* wing disc, not only the source, but also the gradient length *λ*, increases with the growing domain [55]. This gradient behavior can be accounted to the pre-steady-state expansion of the gradient [25]. The combination of an increase in both the gradient amplitude and length results in a gradually rising concentration near the source, a decline at a distance, and a constant concentration at the level of the Dpp readout boundaries for *sal* and *dad*. While this enables the scaling of the Dpp readout positions on the growing domain [25, 56], it precludes the use of the Dpp gradient as timer.

As technological advances enable the quantitative characterization of an increasing number of developmental systems, examples of this novel timer paradigm might be uncovered and the extent to which it reflects the underlying biological process might be clarified. Gradient-based timers present a promising strategy also in synthetic bioengineering efforts to control position and time simultaneously.

## Acknowledgements

We thank Marius Almanstötter, Marcelo Boareto, James Briscoe, José Dias, Johan Ericson, Anna Kicheva for discussions. This work was partially funded by the Swiss National Science Foundation through Sinergia grant CRSII5 170930 to DI.

## Author Contributions

DI conceived the study and discovered the gradient-based timer paradigm. RV and DI developed the theory, analysed the data, and wrote the manuscript. RV created the figures.

## Footnotes

1 Note that a hump can result also in the case of self-enhanced morphogen decay, for which the gradient shape is a shifted power law of the form *C*(*x*) = *C*_0_ (1 + (*x* − *L*_S_)*/mλ*_*m*_)^−*m*^, *m* > 0 [33]. Eq. 9 then reads 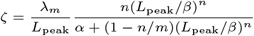.

